# Chronic predation risk induces sex-specific effects in behavior but does not induce long-term oxidative damage

**DOI:** 10.64898/2026.02.23.707534

**Authors:** Michaela M. Rogers, Jennifer K. Hellmann

## Abstract

Predation is a strong environmental and selective pressure that can favour rapid and plastic shifts in behaviour and escape ability to increase an organism’s immediate survival. However, maintaining antipredator responses under repeated predation stress can induce physiological costs to an organism from long-term exposure to elevated cortisol. We know little about how individuals balance this trade-off between short-term survival and longevity, including whether males and females balance this trade-off differently based on life history differences in reproduction, survival, and risk adversity. To assess sex differences in long-term behavioural responses and physiological costs to predation risk, we exposed threespine stickleback (*Gasterosteus aculeatus*) to visual cues of a live rainbow trout (*Oncorhynchus mykiss*) predator twice a week for 14 weeks, then measured stickleback antipredator behaviour and swimming performance 5 months later. To quantify potential long-term costs of behavioural adaptation, we measured relative telomere length as a proxy for long-term oxidative damage. We found strong sex specific effects in behaviour and swim endurance: males, but not females, altered their hiding behaviour and had shorter swim endurance in the first trial, suggesting overall lower activity. Surprisingly, we found no evidence for chronic predation shortening telomere length or hindering growth in body length. Overall, these results suggest that plastic responses can be dictated by the different life-history strategies for males and females, and suggest that individuals can maintain long-term changes in antipredator behaviour without costs to their physiological state.

**Highlights:**

- Chronic predator exposure produced persistent sex differences in space use and swim performance.
- Predator-exposed males altered their hiding strategy and showed reduced swim performance, while females showed no behavioural or performance differences.
- Differences in swim times were restricted to the first trial and all individuals were exhausted by trial 3.
- Relative telomere length and growth in length did not differ between exposed and unexposed individuals.

## INTRODUCTION

Organisms flexibily respond to changes in the environment through phenotypic plasticity, which allows individuals to modify their morphology, physiology, and behaviour in response to environmental cues, even within a single generation (Bonduriansky & Day, 2018). In some cases, this plasticity can be adaptive by allowing individuals to develop phenotypes that match their future environment (Ghalambor et al., 2007). For instance, pugnose shiners (*Notropis anogenus*) exposed to long-term elevated temperatures developed larger gills, an adaptive adjustment to maintain oxygen levels under thermal stress (Potts et al 2021). Similarly, deeper bodies induced by predator exposure in carp enhances their escape locomotor performance (Domenici et al., 2008). However, plasticity can be maladaptive, especially when environmental cues induce changes to an individual’s somatic state (reviewed in Lushchak, 2011). In bulb mites, for example, high nutrition diets cause individuals to develop into larger, more aggressive phenotypes while low nutrition produces smaller and non-aggressive individuals, reflecting growth constraints rather than information about the future environment (Smallegange 2011). Finally, it is possible that this plasticity can come with fitness tradeoffs: an individual may demonstrate adaptive responses to plasticity on a short-term scale (e.g., antipredator responses to predator cues), but there could be trade-offs or negative effects later in life from costs to the individual’s somatic state. For example, grasshoppers exposed to chronic predation escaped faster and jumped further compared to unstressed grasshoppers, but these behavioural modifications in stressed grasshoppers induced a reproductive cost (smaller and less athletic offspring; Hawlena et al., 2011). Understanding when and how plasticity induces fitness costs and benefits is critical for predicting how organisms will respond to both natural and human-induced environmental change.

Predation risk is an ecologically relevant stressor, and plastic changes in prey behaviour have direct links to fitness since the failure to respond appropriately can result in immediate death. However, predicting the fitness outcomes of predator-induced plasticity may be particularly complicated, since early life exposure to predators can induce adaptive antipredator traits (Benard, 2004), but can also induce physiological stress that can reduce an organism’s condition. For example, exposure to predation can heighten antipredator behaviour in future predator encounters, including reduced foraging, spine-erection, predator inspection, freezing, and increased shoaling (Kozak & Boughman, 2012; Landeira-Dabarca et al., 2019). However, changes in antipredator behaviour can also be associated with somatic costs including increased metabolic rates and fewer resources for other necessary somatic functions (Janssens & Stoks, 2014), such as immune function (Rigby & Jokela, 2000). This tradeoff can arise because the accumulation of glucocorticoid hormones (the end product of the hypothalamic-pituitary-adrenal (HPA) stress axis) helps individuals cope with stress, but can also impede long-term growth or induce oxidative stress (Nicolaides et al., 2015). Consequently, the maintenance of homeostasis under high predation conditions can induce a trade-off between long-term health and shorter-term survival (Kim et al., 2019; Nicolaides et al., 2015), especially when exposure persists over longer timescales. However, despite this recognition, our understanding of how predation early in life simultaneously influences behaviour and physiology later in life remains limited.

Individuals may differ in the ways in which they manage this trade-off between short-term survival and long-term phenotypic effects. In particular, a growing body of work underscores the importance of considering sex differences in response to stressors, including predation risk (Hellmann et al., 2020). Males and females frequently differ in their life-history strategies, reproductive investments, and exposure to predation which leads to unequal selective pressures that shape antipredator phenotypes and the fitness benefits associated with expressing them (Andersson 1994, Gosline and Rodd 2008). This could result in several different outcomes. First, one sex might be more vulnerable to predation compared to the other, such that the benefits of increasing short term survival with antipredator behaviours outweigh the cost to longevity through somatic costs. In contrast, the less susceptible sex may be more likely to ignore the predator cues in order to avoid long-term costs. For example, predation risk experienced by cichlids early in life caused increased antipredator morphologies in males; but not females, possibly because males are more vulnerable to predation (Meuthen et al., 2018). Second, both males and females could mount plastic responses to predation risk, but the sexes could differ in the extent to which those phenotypic changes result in adaptive traits versus long term somatic costs. For example, predation risk in snails suppressed growth only in males, while risk eliminated the positive relationship between size and fecundity in females, without affecting overall fecundity (Donelan & Trussell, 2020). This pattern suggests that predation risk altered the condition-dependent allocation of reproductive resources in females, with females displaying weaker antipredator responses in order to maintain energy needed for reproduction. Understanding why these sex differences arise is important for knowing how responses to predation may be driven by differences in individual ability to mount and sustain stress responses.

Stickleback are an excellent system for exploring sex-differences in long-term responses to predation risk. Antipredator responses in stickleback are under selection (Giles & Huntingford, 1984; Wund et al., 2015), with longer spines and more armor in high predation populations compared to low predation populations (Marchinko, 2009). Within a population, males and females also differ in vulnerability and ability to respond to predators. For example, while females are free-swimming and tend to form shoaling groups, males typically stay within a territory while nesting in limnetic areas (FitzGerald, 1993). Males also have more conspicuous reproductive coloration and courtship behaviour than females (Candolin, 1999, 2000). This suggests that optimal responses to predation may differ between stickleback males and females, such that females may benefit from increased swim endurance while males may benefit more from increased hiding.

To investigate the long-term effects of predation on behavioural and physiological traits, we exposed juvenile male and female threespined stickleback (*Gasterosteus aculeatus)* to visual cues of a rainbow trout *(Oncorhynchus mykiss)* for 14 weeks. One month after the last exposure, we re-exposed individuals for a week and gave them a 5 month break before we tested their antipredator behaviour, swim endurance, and took tissue samples to measure relative telomere length. While trout are a novel predator for this population of sticklebacks, previous studies show threespined stickleback respond to visual cues of rainbow trout predators and increase their antipredator behaviour after exposure to trout cues (Messler et al., 2007; Rogers & Hellmann, 2025). Therefore, if predation exposure in early life adaptively primes individuals for predation risk later in life, then we expect increased changes in behaviour, longer swim endurance and increased growth in predator-exposed individuals. In constrast, if experience with predation risk in early life has maladaptive long-term effects, then we expect to see shorter swim endurance, shorter telomeres, and reduced growth in predator-exposed individuals. It is also possible that we see fitness trade-offs: predation in early-life could better prepare individuals to cope with future predation through changes in behaviour (e.g., increased swim endurance, heightened antipredator behavor), but compounded stress may come at a cost of reduced physiological fitness (e.g., shorter telomeres). Because male stickleback are generally more vulnerable to predation than females, we further predict that males may exhibit stronger antipredator behavioural responses following early-life predator exposure, but potentially incur greater physiological costs such as shorter telomeres.

## METHODS

### Sourcing and Housing

The fish used in this experiment were lab-reared, F1 descendents of adult threespined stickleback *(Gasterosteus aculeatus)* collected from Putah Creek (CA, USA) in June 2021. Prior to the experiment, we housed individuals on a recirculating system in groups of 10–12 fish per tank to mimic shoaling conditions in the wild. We maintained the system on a summer photoperiod schedule (16L:8D) at 20 ± 1°C and fish were fed once a day ad libitum with a mixture of frozen bloodworms (*Chironomus* spp.), brine shrimp (*Artemia* spp.), Mysis shrimp and Cyclop-eeze. We obtained rainbow trout *(Oncorhynchus mykiss)* from Freshwater Farms of Ohio in June 2021. Rainbow trout are present in Putah Creek, but they are not present in Beaver Pond (J.K.H. & A. Bell, personal observation), which is an isolated section of Putah Creek where we collected the stickleback. However, previous work on this population showed within- and transgenerational responses to visual cues of rainbow trout (Hellmann & Rogers, 2024; Rogers & Hellmann, 2025).

### Exposure regime

In May 2022, we moved n = 83 individuals in groups of 10-11 into 8 experimental 37.9 L (53 L × 33 W × 25 H cm) stand-alone tanks. We mixed groups to have 2-3 individuals from the same clutch and at least one female and one male from the same clutch (for a total of n = 27 genetically distinct clutches). We initially marked individuals by clutch with an injection of elastomer dye. The tanks contained an airstone, gravel, and two fake plants. Between each experimental tank, we placed another 37.9 L tank filled with freshwater and containing gravel and two sponge filters. Depending on the visual predator treatment, the tank was either left empty or contained one live rainbow trout.

After at least 7 days in the tank to habituate, we exposed stickleback twice a week for 14 weeks to a 2-hour visual cue of a rainbow trout. All experimental tanks were separated visually from each other with a removeable opaque divider. We exposed fish randomly twice a week, at a random time between 9:00 and 16:00. In the control treatment, we turned off the airstones in the tanks and removed the divider to show a tank filled with only water. In the visual predator treatment, we turned off the airstones in the tanks and removed the divider to show a live trout in the adjacent tank. To reinforce the visual cue of the predator, we fed the trout two feeder guppies *(Poecilia reticulata)* at the beginning of the 2□h exposure. After 2 h, we replaced the dividers and turned on the airstones. A schematic of the experimental timeline and exposure tank setup is shown in Figure 1. After the 14 week exposure, we measured length and mass for all individuals (“initial” measurement; n = 59). One month after the last exposure, we exposed individuals for one more week (two exposures of a 2-hour visual cue) to reinforce the cue; there were no exposures for 5 months prior to beginning assays. Starting in March of 2023, individuals went through an open field assay and a swim endurance assay. 14 individuals (n = 10 control, n = 4 predator-exposed) died during the 9 months from when exposures started to when the break ended. Additionally, we didn’t use n = 10 of the individuals because of deformities that would have affected their behaviour and swim performance.

**Figure 1.**
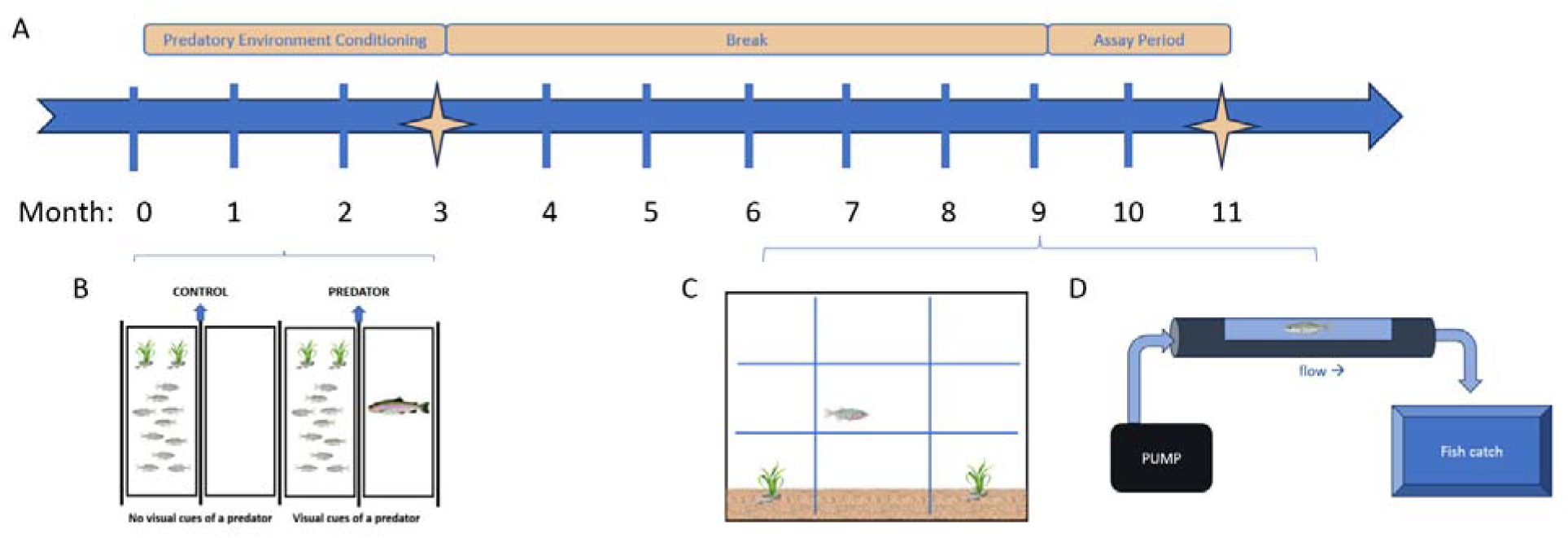
Overview of the experimental timeline (A). The tank setup for the exposures are shown in (B), a side view of the open field assay is shown in (C), and a schematic of the swim endurance assay is shown in (D). The stars at months 3 and 11 mark the two measurement timepoints for individual standard length and mass.

### Behavioural Testing

From March – April 2023, we tested individuals in an open field assay to measure behaviour before and after exposure to a model trout predator (Figure 1C). 24-hours before the assay, we added another mark using elastomer dye to mark each individual and moved them to an individual holding tank. The behavioural arena was a 37.9 L tank containing gravel and two fake plants and was divided into 9 rectangular sections on the front of the glass. To begin, we released the focal fish from a cup into the tank and allowed them to acclimate freely in the tank for 15 minutes to habituate. After 15 minutes, we scored the amount of time in each section (including the amount of time spent in the bottom third of the tank as a proxy for hiding), time spent near the plants, and time frozen (as a proxy for perceived risk) using BORIS tracking software (Friard & Gamba, 2016) for 5 minutes (‘baseline’ behaviour). Then we chased the fish for 15s with a 10 cm model of a rainbow trout. 15s after the simulated predator attack, we resumed scoring the same behaviours for 5 minutes (‘exposed’ behaviour). Then we placed them back in their individual holding tanks before the next assay.

24-hours after the behavioural assay, we ran individuals through a swim endurance assay (Figure 1D). We built the swim assay chamber with a pump to push water through a flow chamber with a viewing window, which flowed into a net box to catch the fish. We set the flow at a consistent rate of 7.7 L/min when the pump was turned on. Individuals acclimated for 5 minutes in the viewing chamber with the pump turned off, then we turned on the water flow and we recorded time spent swimming against the flow for up to 15 minutes or until they were caught in the net box due to exhaustion. We then placed the fish back in the viewing chamber for 5 minutes before starting trial 2. This process was repeated to achieve 3 consecutive trials with 5 minutes of acclimation in the viewing chamber in between each trial. 48-hours following the swim assay, we again measured individuals for length and mass (“final” measurement) and then dissected individuals for sexing by identifying their reproductive organs and removed a piece of muscle adjacent to the caudal fin to collect DNA samples for relative telomere length. In total, we used n = 28 control individuals (n = 10 males and n = 18 females) and n = 31 predator-exposed individuals (n = 7 males, n = 24 females).

### Analysis of Telomere Length

We measured telomere length in muscle DNA samples (see above) by modifying the qPCR method developed by Cawthon (2002) and validated in threespine stickleback by Kim et al. (2019). We used the same control gene as Kim et al. (2019), the threespine stickleback-specific glyceraldehyde-3-phosphate dehydrogenase gene (GAPDH). For each sample, we diluted the DNA to 10 ng for both telomere and GAPDH reactions. The primer sequences (written 5′→3′) were: Telomere forward (CGGTTTGTTTGGGTTTGGGTTTGGG TTTGGGTTTGGGTT) and Telomere reverse (GGCTTGCCTTACC CTTACCCTTACCCTTACCCTTACCCT); threespine stickleback-specific GAPDH forward (GAGACGTGACCATTGAGGGG) and reverse (TGTGCGGTGGGCTTTATGAT). We validated the concentrations of primers to be used in the qPCR assay using standard curves. Telomere primers were used at a concentration of 300 nM for the forward and 500 nM for the reverse. GAPDH primers were used at a concentration of 500 nM for the forward and 700 nM for the reverse. DNA samples and primers were mixed with 10 μl of PowerTrack SYBR Master Mix (Fisher Scientific) for a total volume of 20 μl. Telomere and GAPDH reactions were performed on separate plates. We used the following qPCR conditions: telomere 5 min at 95°C, followed by 30 cycles of 15 s at 95°C, 60 s at 60°C, followed by 15 s at 95°C, 60 s at 60°C, then 15 s at 95°C; and GAPDH 5 min at 95°C, followed by 40 cycles of 15 s at 95°C, 60 s at 60°C, followed by 15 s at 95°C, 60 s at 60°C, then 15 s at 95°C. For each 96-well plate, we included a DNA reference sample from threespine stickleback , created by mixing DNA extracted from multiple individuals and diluted to 10 ng, and a “no template control”, to check that the reagents were not contaminated. We calculated relative telomere length (RTL) by determining the ratio (T/S) of telomere repeat copy number (T) to a single control gene copy number (S), relative to a reference sample.

### Statistical Analysis

To determine mean differences in individual behaviours in the open field and swim endurance assays, we ran MCMC generalized linear mixed models (R package MCMCglmm; Hadfield, 2010). All models had 200,000 iterations, a burn-in of 3,000 iterations, thin = 3 and Poisson distributions. For the open field assay, we ran models for time spent in the bottom third of the tank, time spent within one body length of either plant, and time frozen. We tested fixed effects of treatment, sex, observation period, and length at the time of assays. We included random effects of observer, clutch identity and individual identity to account for paired data. For all models, we included interactions between treatment, sex, and observation period, then removed any nonsignificant interactions to arrive at the final models.

For the swim endurance assay, we were specifically interested in interactions between treatment, sex, and trial number to assess factors contributing to differences in swim time across the three trials, with random effects of fish identity and clutch identity. Due to a 3-way interaction between treatment, sex, and trial number in models with all 3 trials combined, we ran separate models for each of the three trials with fixed effects of treatment, sex, and final length, and random effects of original exposure tank and clutch. We again tested for interactions between treatment and sex and removed nonsignificant interactions as necessary.

For RTL, we ran MCMC glmms with Gaussian distributions, including fixed effects of treatment, sex, and final body length, with clutch and qPCR plate as random effects. Model outputs were similar if we controlled for mass instead of length. Additionally, mass is less informative due to females becoming gravid throughout the experiment, therefore we report results with length below only. We removed six samples from the analysis due to poor replications in the qPCR assay ending up with a total of n = 53 samples (n = 26 control, n = 27 predator-exposed).

To determine what traits may have had an effect on growth, we took two measurements of length and mass, once after the 14 weeks of exposure (“initial”), and once after assays and prior to dissection to collect tissue for measuring RTL (“final”). We subtracted initial from final to calculate growth for both length and mass. We ran MCMC models with 200,000 iterations, a burn-in of 3,000 iterations, thin = 3 and Gaussian distributions with a weak prior. To predict growth in length across the experiment, we included fixed effects of treatment, sex, and initial length, with a random effect of clutch. To predict growth in mass across the experiment, we included fixed effects of treatment, sex, and initial mass, with a random effect of clutch.

### Ethical Note

All methods were approved by the Institutional Animal Care and Use Committee of The Ohio State University (protocol ID 2023A00000055), including the use of live and model predators, and adhere to the guidelines set forth by the Animal Behaviour Society and the Association for the Study of Animal Behaviour. In all parts of this study, care was taken to minimize stress to the study animals. To identify stickleback individually, we marked them with elastomer dye and did so efficiently to minimize time out of the water; there were no ill effects or mortality due to the injection. The live rainbow trout (*Oncorhynchus mykiss*) predators used for exposures were kept in individual tanks with an airstone and plants for refuge and were returned to group housing in a 208L tank (123.8 L × 35.6 W × 54 H cm) after exposures concluded. The predators were fed daily with pellets, except during exposures when they were fed 2 live feeder guppies. Guppies were eaten immediately upon being dropped in with the live predator and were necessary to reinforce the visual cue, which would otherwise be nonlethal for the sticklebacks. To reduce the use of live predators and use the minimum amount of stress necessary to the study organism, a model predator of a rainbow trout was used in the behavioural assay to be able to quickly chase the focal fish. When moving sticklebacks between holding tanks and assay chambers, time out of the water was minimized as much as possible by catching fish with a net and immediately putting them in a cup of water to transport them. Between assays, stickleback were able to recover overnight in their holding tanks containing gravel and two plants. Following the last assay, all stickleback were euthanized swiftly by decapitation for the purpose of obtaining tissue for relative telomere length and sexing. In total, 14 of the initial 83 experimental fish died before the end of the experiment. Wild stickleback in our population typically die after about a year, so some natural mortality was expected across the length of our experiment.

## RESULTS

To understand long-term behavioural and physiological consequences of predation risk, we exposed individuals to novel trout predators for 14 weeks. One month after the last exposure, we exposed them for one more week before giving them a 5 month break. Following the break, we ran individuals through an open field assay and a swim endurance assay and collected muscle tissue to measure relative telomere length.

### Open Field Assay

We exposed individuals to long-term visual cues of a rainbow trout predator and measured their behaviour in an open field assay pre- and post-exposure. We found a significant interaction of treatment and sex for time spent in the bottom third of the tank (MCMC GLMM, 95% CI in brackets here and below; treatment*sex: [0.30, 4.27], p = 0.02, Table 1, Figure 2a). When subsetting by sex, we did not find a significant effect of treatment in males ([-0.96, 3.87], p = 0.20) or females ([-1.20, 0.91], p = 0.77). However, when we subset by treatment, we found that predator exposed males spent significantly more time in the bottom third of the tank (in both the pre- and post-periods), than predator-exposed females ([0.04, 3.51], p = 0.04) but this sex difference was not seen between control males and females ([-2.72, 0.32], p = 0.11). We also found a significant treatment by sex interaction for time spent hiding in the plant (treatment*sex: [0.55, 6.64], p = 0.02, Table 1, Figure 2b). Subsetting again by sex, we found that predator-exposed males spent less time hiding than control males (treatment: [2.30, 13.66], p = 0.005) and no change in hiding for females (treatment: [-1.31, 1.44], p = 0.96). This pattern was also reflected when we subset by treatment. When subsetting by treatment, we saw the opposite pattern than time spent in the bottom third of the tank: control males spent less time than control females in the plant ([-7.96, -0.50], p = 0.01), while there were no sex differences in the predator treatment ([-2.71, 2.06], p = 0.72).

**Table 1:** Results of MCMCglmm models testing predictors of time spent in the bottom third of the tank and time spent hiding in the plant in the behavioural assay, as well as swim time in the swim endurance assay. We tested for potential interactions between treatment, sex, and trial when possible. We removed any interactions that were not statistically significant. Bold values indicate statistically significant *p*-values.

| Time in the bottom third |  |  |  |
| --- | --- | --- | --- |
|  | Mean | 95% CI (L, U) | *p* |
| Treatment | 3.72 | (-1.38, 0.77) | 0.56 |
| Sex | -0.32 | (-2.33, 0.50) | 0.21 |
| Observation period | -0.89 | (0.49, 1.99) | **0.002** |
| Length | 1.24 | (-0.16, 0.12) | 0.77 |
| Treatment * Sex | -0.02 | (0.30, 4.27) | **0.02** |
| Time near plant |  |  |  |
|  | Mean | 95% CI (L, U) | *p* |
| Treatment | 0.50 | (-1.12, 2.07) | 0.53 |
| Sex | -2.90 | (-5.13, -0.68) | **0.008** |
| Observation period | 0.55 | (-0.06, 1.17) | 0.08 |
| Length | 0.10 | (-0.11, 0.31) | 0.31 |
| Treatment * Sex | 3.58 | (0.56, 6.64) | **0.02** |
| Swim time |  |  |  |
|  | Mean | 95% CI (L, U) | *p* |
| Treatment | 0.27 | (-0.77, 1.28) | 0.60 |
| Sex | 2.59 | (1.29, 3.91) | **<0.001** |
| Trial 2 | 0.46 | (-0.57, 1.46) | 0.37 |
| Trial 3 | -0.58 | (-1.58, 0.44) | 0.26 |
| Treatment * Sex | -3.89 | (-5.77, -1.95) | **<0.001** |
| Treatment * Trial 2 | -0.32 | (-1.63, 1.05) | 0.63 |
| Treatment * Trial 3 | -0.44 | (-1.80, 0.88) | 0.52 |
| Sex * Trial 2 | -1.53 | (-3.20, 0.15) | 0.07 |
| Sex * Trial 3 | -2.98 | (-4.66, -1.27) | **<0.001** |
| Treatment * Sex * Trial 2 | 1.90 | (-0.63, 4.37) | 0.14 |
| Treatment * Sex * Trial 3 | 3.68 | (1.19, 6.23) | **0.003** |

**Figure 2a.**
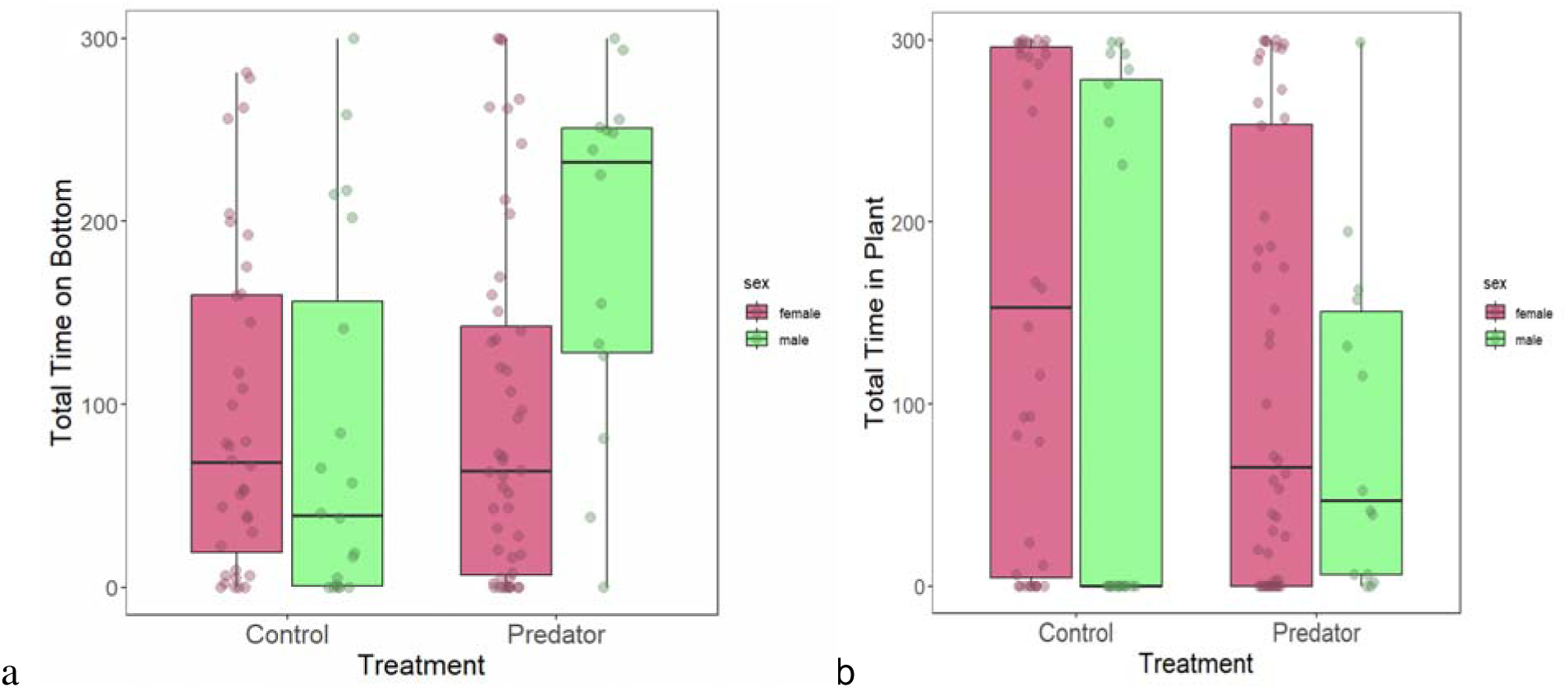
Combined^1^ time spent by females (pink) and males (green) in the bottom third of the tank (data are median with interquartile range). Predator-exposed males spent more time on the bottom compared to predator-exposed females, and this difference was not seen between control sexes. 2b. Combined^1^ time spent in the plant by females and males. Males typically spent less time in the plants regardless of treatment or exposure-period. ^1^Combined values mean there are two datapoints per individual in each boxplot, one for time in the pre-period and one for time in the post-period.

Time frozen was measured in both the pre- and post-exposure periods. All individuals, regardless of treatment or sex, froze significantly more after the predator stimulus compared to before, confirming that individuals were behaviourally responding to the simulated predator attack (observation period: [2.95, 6.72], p < 0.001. We found no effect of treatment ([-2.24, 1.68], p = 0.81), sex ([-2.17, 2.24], p = 0.98), or length ([-0.43, 0.13], p = 0.32) on time frozen.

### Swim Assay

We ran individuals through 3 consecutive swim endurance trials with 5 minutes of acclimation prior to each trial. We found a significant 3-way interaction between treatment, sex, and trial (3-way interaction: [1.19, 6.23], p = 0.004; Table 1, Figure 3).

**Figure 3.**
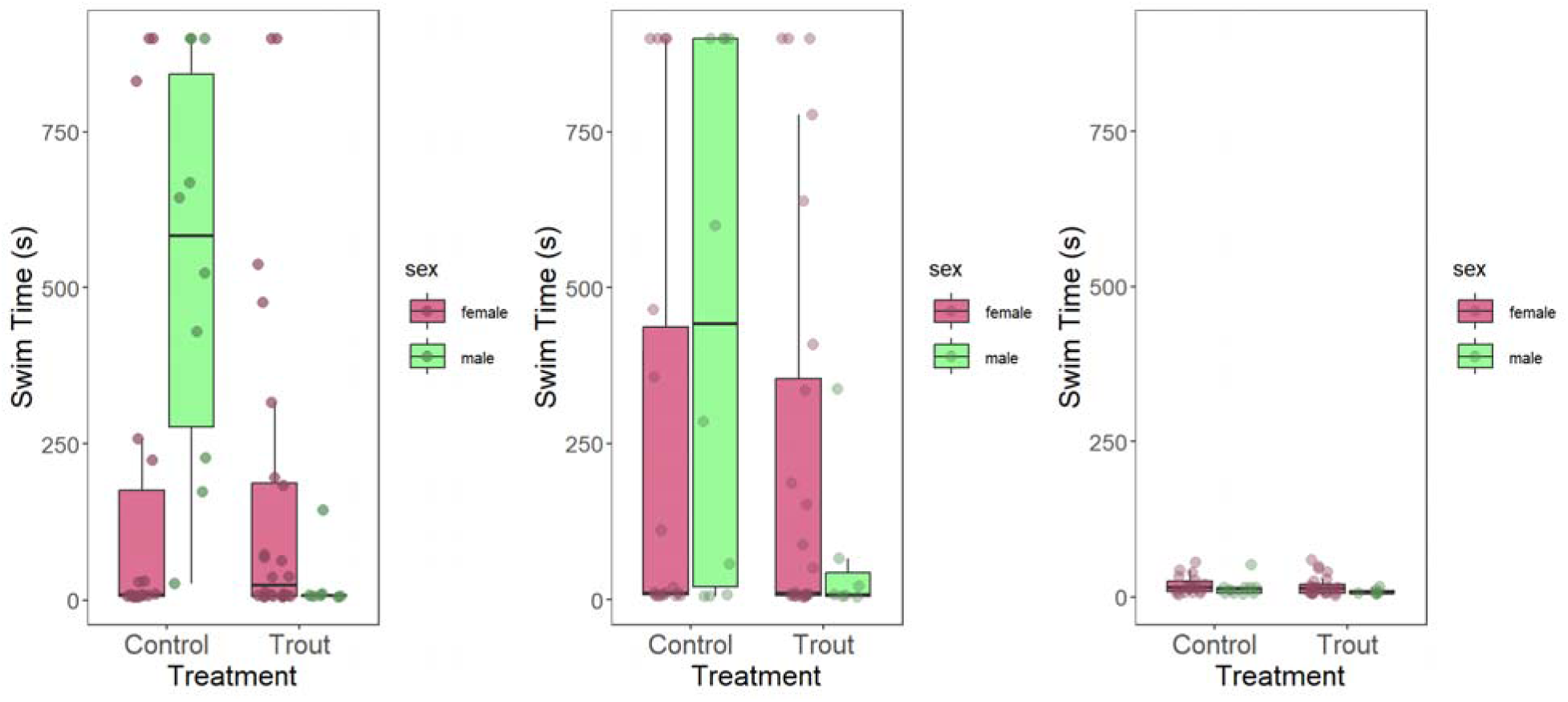
Swim time across all 3 trials (from left to right) by treatment. Pink boxplots are female and green boxplots are males (data are median with interquartile range, dots correspond to individuals). In trial 1, unexposed males swam longer than predator-exposed males, while there was a trend for predator-exposed females to swim longer than unexposed females. This interaction between treatment and sex did not persist past trial 1.

*Post-hoc* analyses for individual trials revealed a treatment by sex interaction in trial 1 (treatment*sex: [-5.78, -1.60], p < 0.001). Specifically, unexposed males swam longer than predator-exposed males in trial 1 ([-5.30, -2.13], p = 0.001), while there was no difference between swim times for predator-exposed females and unexposed females ([-1.00, 1.86], p = 0.55). By trial 2, there were no differences between swim times across treatments or sexes (treatment: [-1.45, 1.12], p = 0.82; sex: [-0.43, 2.34], p = 0.18). In trial 3, we did not find a significant effect of treatment ([-0.82, 0.24], p = 0.27), however, there was a marginal trend for females to swim longer than males ([-0.96, 0.04], p = 0.08). Swim times in trial 3 were drastically shorter for all individuals compared to other trials, with no individuals swimming longer than 60 seconds.

### Relative Telomere Length (RTL)

We found that sex marginally predicted telomere length (sex: [-0.32, 0.01], p = 0.06), with males tending to have shorter telomeres than females (Figure 4a). There was no effect of treatment ([-0.22, 0.06], p = 0.25) or final length (−0.03, 0.01], p = 0.57) on relative telomere length.

**Figure 4a.**
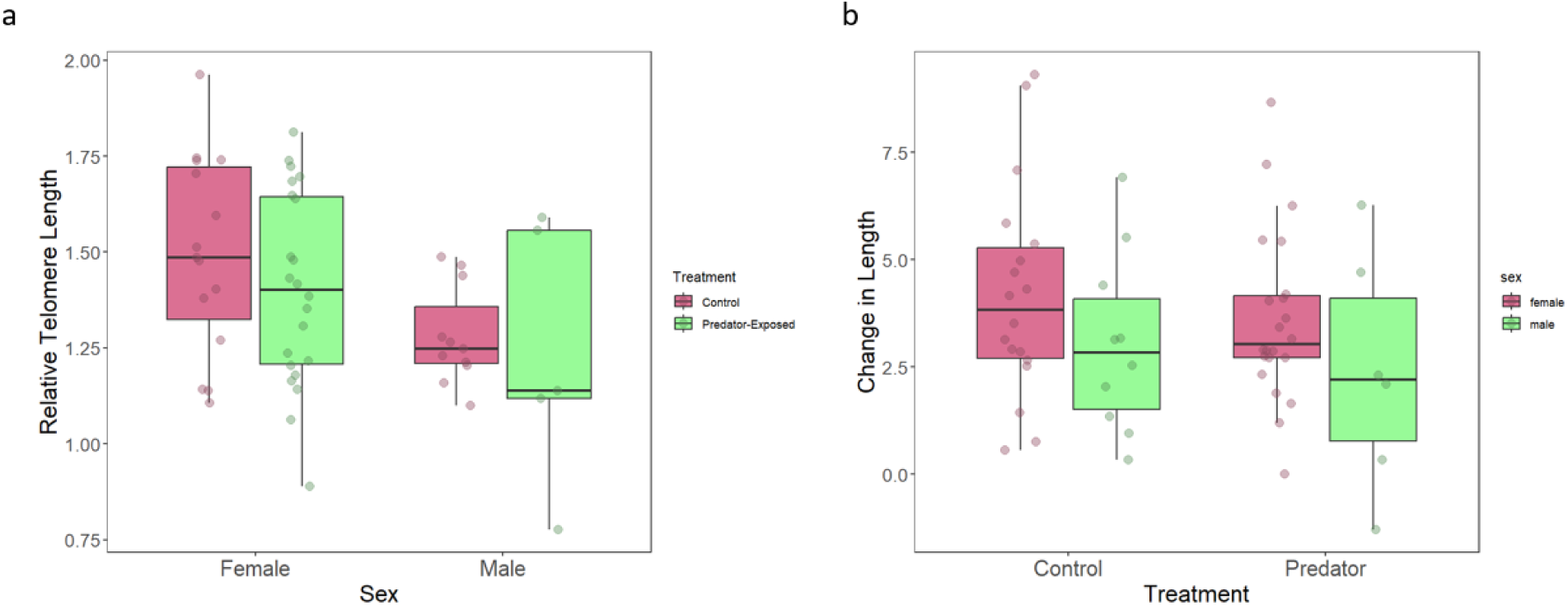
Relative telomere length by sex with treatment (data are median with interquartile range, dots correspond to individuals). 4b. Change in length for unexposed and predator-exposed males and females across the two measurement timepoints. All individuals had a positive change in growth except for one predator-exposed male.

### Growth

We measured individuals at two timepoints approximately 7 months apart, first at the time of exposures (initial) and again at the time of assays (final) to calculate growth from the difference between the two measurements. We did not find a significant effect of treatment on change in length, although predator-exposed individuals tended to grow less during that time (treatment: [-2.26, 0.10], p = 0.09). However, we did find a significant effect of sex and initial length on change in length (Figure 4b). Specifially, we found that males were bigger than females at the time of exposures (initially; sex: [-4.22, -0.36], p = 0.02), and that males grew faster than females during the break (sex: [-319, -0.54], p = 0.007). Therefore, individuals initially bigger at the time of exposures were also bigger by the end at the time of assays (length: [-0.41, 0.06], p = 0.01). It is important to note that our sample size for males in the predator treatment was n = 5; individuals were selected for the experiment prior to sexual maturity, which did not allow us to ensure even sex ratios.

## DISCUSSION

Plasticity in response to environmental cues can be adaptive by priming individuals with information about their future environment, but it can induce lasting physiological costs that constrain performance, growth, and longevity later in life. We first exposed individuals to long-term predation before giving them an extended recovery period, then measured behaviour, growth, and relative telomere length. We found sex- and treatment-specific responses in behaviour and swim endurance: males exposed to predation risk spent more time in the bottom of the tank during an open field assay and swam for shorter periods of time during the swimming trials, while females showed no differences in behaviour or swimming performance. However, we found no evidence that predation exposure induced changes in growth or telomere length for neither males nor females. Our results reveal that early-life predation risk acts to shape behavioural phenotypes in sex-specific ways, but does not appear to be a chronic stressor that degrades physiological traits over time.

Early-life predation risk had the strongest effect on behaviour, particularly through sex by treatment interactions in space-use. Predator-exposed males spent more time in the bottom third of the tank, while control males used the plants for hiding following the model predator stimulus in the open field assay. In contrast, predator-exposed females showed no differences in behaviour compared to the control females. Males spending more time in the bottom third could reflect lower activity levels in predator-exposed males compared to control males. Similar to previous studies that have seen reduced activity in response to cues of predation (Lacasse & Aubin-Horth, 2012; Stein & Bell, 2014), it could reflect a consequence of predation being stressful. Alternatively, staying in the bottom third of the tank could reflect a hiding behaviour to avoid the open water column. Typically, non-shoaling fish species do not use evasive tactics to avoid predators, and often freeze to reduce movement and avoid detection by predators (Lu et al 2025). Therefore, males with prior predator experience potentially opted to stay close to the gravel instead of moving to one of the two plants during the assay. Overall, we see that prior predator experience changed how males hide, rather than how much they hide, facilitated by having multiple measures of hiding in this experiment. These patterns suggest that predation risk may provide information that reshapes how individuals assess and manage risk, rather than simply suppressing or increasing activity across treatment groups (Stein & Hoke, 2022). This also suggests that it is important to consider the type of response in addition to the magnitude of response, in the case that individuals trade off between different antipredator responses.

For swim performance, we expected that individuals with early life predation experience would swim longer in the swim endurance assay compared to unexposed individuals. Previous studies have found plastic morphological changes in response to predation risk increase escape speed in the presence of live predators (Domenici et al., 2008; McPeek et al., 1996; Teplitsky et al., 2005). However, we only saw differences among treatment groups in trial 1: predator-exposed males had shorter swim times than unexposed males, while females maintained similar swim times across treatments. By trials 2 and 3, there were no differences among treatments or sexes. Differences in male swim times in trial 1 could reflect reduced endurance because of trade-offs with physiological capacity. For example, predator-exposed damselflies had reduced escape speed because of an increase in oxidative damage from chronic predation stress (Janssens & Stoks, 2014). However, the absence of significant treatment effects on physiological traits, including male growth and telomere length, suggests that the predation stress did not impose a lasting physiological cost to male somatic state. Futher, the fact that treatment differences in swimming ability did not persist past the first trial suggests that predator-exposed individuals do not have lower endurance. Instead, reduced swim performance in predator-exposed males could be from a lack of motivation to participate in the assay. This is consistent with the fact that predator-exposed males spent the most time on the bottom of the tank in the open field assay. Together, increased time on the bottom and shorter swim times suggest that predator-exposed males end up with a generally lower activity level following chronic predator exposure.

The sex-specific nature of these behavioural and performance responses is consistent with known differences in stickleback life history, ecology, and predation vulnerability. Males are typically more conspicuous than females because they hold nesting territories and conduct courtship displays for females (FitzGerald, 1993). They also develop bright reproductive coloration, which increases exposure to predators (Candolin, 2000). In contrast, females are typically free-swimming and create shoals, reflecting the benefit of predator-exposed females maintaining swim endurance compared to unexposed females. These observed patterns suggest that early-life predation risk may shift males towards more conservative, risk-adverse strategies; this aligns with evidence that males across other taxa are usually more vulnerable to predation (reviewed in Magnhagen, 1991; Meuthen et al., 2018; White et al., 2022) and previous evidence in stickleback show that males are more responsive to direct cues of predation risk (Hellmann et al., 2020; Herczeg et al., 2015). In contrast, there may be greater selection for females to maintain their behaviour and swim performance, as slower escape speed can put individuals at a disadvantage in a shoal (Frommen et al., 2012). These sex-specific responses likely indicate different optimal phenotypes for males and females under predation which are shaped by their different reproductive roles and ecological niches. Assessing sex differences is important for understanding how plasticity evolves under environmental stress because sex-specific roles can determine which traits are under selection and reveal the different constrants that males and females face in altering their behavioural and performance.

Despite evident sex effects of early-life predation on behaviour and swim performance, we found no strong evidence that predator exposure hindered growth or reduced telomere length. Instead, we found effects of sex independent of treatment, with males growing more and tending to have shorter telomeres compared to females. The lack of an effect of treatment could either suggest that the physiological stress from over 3 months of exposure was insufficient to impose the cost of telomere shortening, or, that individuals may have been able to buffer telomere loss during the break. It is unclear if we would have found a stronger effect of treatment on growth if we had measured individuals for the initial measurement before the 14 week exposure rather than after. Overall, the lack of connection between behavioural responses and growth and telomere length suggest that early-life predation in this system largely affects plastic antipredator responses rather than imposing long-term costs. This suggests that plastic responses may not be costly to produce and can inform predictions about survival, reproduction, and population dynamics under changing environments.

In conclusion, our results support the prediction that individuals did not have uniformly adaptive or maladaptive traits and instead had a suite of behaviours that was expressed plastically, independently of physiological condition or somatic state. Knowing that behavioural plasticity and physiological condition are not necessarily inversely related may guide future directions to examine how predator exposure across life stages shapes long-term fitness, reproductive effort, and survival against predators. For example, a meta-analysis found that reproductive strategies are associated with sex differences in resistance to oxidative damage (Costantini, 2018). This suggests that behaviour should be measured together with other somatic traits to be able to to fully assess the trade-offs that individuals face when they are dealing with stressors.

## ACKNOWLEDGEMENTS

This work was supported by the University of Dayton, the Ohio State University, and by National Science Foundation Grant # IOS 2446322. We thank Emerson Amy, Amy Friemoth, and Sarah Metz for their help with smoothly and efficiently running the behavior and swim performance assays. We would also like to thank Julie Reynolds and Elizabeth George for their help with the use of the qPCR machine and measuring relative telomere length. Finally, we thank the members of the Hellmann Lab for their comments on previous versions of this manuscript.

## Notes

### Competing Interest Statement

The authors have declared no competing interest.

